# Multi-modal brain magnetic resonance imaging database covering marmosets with a wide age range

**DOI:** 10.1101/2022.09.21.508952

**Authors:** Junichi Hata, Ken Nakae, Hiromichi Tsukada, Alexander Woodward, Yawara Haga, Mayu Iida, Akiko Uematsu, Fumiko Seki, Noritaka Ichinohe, Rui Gong, Takaaki Kaneko, Daisuke Yoshimaru, Akiya Watakabe, Hiroshi Abe, Toshiki Tani, Henrik Skibbe, Masahide Maeda, Frederic Papazian, Kei Hagiya, Noriyuki Kishi, Tomomi Shimogori, Tetsuo Yamamori, Hirotaka James Okano, Hideyuki Okano

## Abstract

Magnetic resonance imaging (MRI) is a noninvasive neuroimaging method beneficial for the identification of normal developmental and aging processes and data sharing. Marmosets have a relatively shorter life expectancy (approximately 10 years) than other primates, including, humans because they grow and age faster. Hence, the common marmoset model is effective in aging research. The current study investigated the aging process of the marmoset brain and provided an open MRI database on marmosets with a wide age range. The Brain/MINDS Marmoset Brain MRI Dataset contains brain MRI information on 216 marmosets aged between 1 and 10 years. During its release date, it is the largest public dataset worldwide. Further, it comprises multi contrast MRI images. In addition, 91 of 216 animals have corresponding ex vivo high-resolution MRI datasets. Our MRI database, which is available at the Brain/MINDS Data portal might help understand the effects of different factors, such as age, sex, body size, and fixation, on the brain. Moreover, it can contribute to and accelerate brain science studies worldwide.

## Background & Summary

Aging is associated with the deterioration of brain function, including cognitive function. Brain magnetic resonance imaging (MRI) databases on humans of various age groups have been published. The integration of these databases has revealed changes in average brain volume with age (e.g., Brain Chart). Changes in the normal aging brain can result in the development of conditions, such as Alzheimer’s and Parkinson’s disease, thereby leading to a better understanding of aging and diseases^1^.

Marmoset (*Callithrix jacchus*), a non-human primate, is useful in aging models^2,3,4^. Hence, it has received significant attention. Compared with rodents, marmosets have a relatively more similar brain structure to humans. Therefore, they are a more suitable preclinical animal model for diseases established via drug administration^6^ and/or genetic manipulation^7^. Furthermore, these animals have a relatively shorter life expectancy (approximately 10 years in captivity) than other primates, including humans. Thus, the aging process is easier to follow. Previous studies have investigated and published MRI datasets on developmental stages or early adulthood among normal marmosets. Seki et al. (2017) provided open-access structural brain datasets (T1-weighted [T1w] and T2-weighted [T2w]; 0–2 years old)^8^. Liu et al. (2020, 2021) published multi-modal, high-quality datasets (T1w, T2w, diffusion-weighted imaging [DWI], awake, anesthetized resting-state functional MRI [rsfMRI]; 3–4 years)^9,10^ However, it has not yet covered the whole life span, including middle (5–7 years) and late (8-10 years) adulthood. To identify the aging process of the marmoset brain, we provided an open-accessible MRI dataset covering the whole life stage of the common marmoset.

The current database contains multi-modal brain MRI datasets on 216 marmosets aged 1–10 years in both in vivo and postmortem studies. The male-to-female ratio in the dataset was 4:6, and data on the weight of animals in each measurement were also available. Hence, the effects of sex and body size on the brain could be examined. All 216 datasets contained T1w and T2w images, and 126 datasets have two-shell in vivo DWI images. Moreover, 31 datasets have 10-min anesthetized rsfMRI images, and 3 have 20-min awake rsfMRI images (total: 60 min). In addition, 91 animals underwent postmortem scans with T2w and DWI images. To the best of our knowledge, this MRI database is the largest to date. Multi-modal datasets including higher-resolution postmortem data could provide not only detailed reliable structural and functional data and its connectivity information but also essential details on the effects of aging on the brain. This openly-accessible database may significantly contribute to the brain science community.

This database can deepen our understanding of the effects of several factors affecting the brain. For example, the volume of gray matter was found to decrease with age. In addition, the DWI structural connectivity showed that most of the connectivity peaked at approximately 3–4 years of age. Moreover, the strength of the connections was significantly lower in the anesthesia state than in the awake state. These findings are consistent with known human facts, and they support the validity of the marmoset developmental and aging model.

## Methods

### Animals in vivo

This study included 216 healthy common marmosets (88 male and 128 female, mean weight: 357.1 ± 60.2 g) aged 0.8–10.3 (mean: 4.34 ± 2.56) years. The common marmosets were anesthetized, and their heads were immobilized before imaging. The in vivo MRI scan was conducted while maintaining each animal in the supine position on an imaging stretcher under anesthesia with 2.0% isoflurane (Abbott Laboratories, Abbott Park, IL, the USA) in an oxygen and air mixture. Heart rate, SpO2, and rectal temperature were regularly monitored during imaging to manage the physical condition of animals. This study was approved by the Animal Experiment Committees of RIKEN Center for Brain Science (CBS) and conducted in accordance with the Guidelines for Conducting Animal Experiments of RIKEN CBS.

### Image acquisition in vivo

The marmoset brain MRI dataset (NA216) contains multi-modal neuroimaging data comprising in vivo T1w, T2w, DWI, and rsfMRI images. MRI was performed using a 9.4-T BioSpec 94/30 unit (Bruker Optik GmbH, Ettlingen, Germany) and a transmitting and receiving coil with an 86-mm inner diameter. For T1w imaging, a magnetization-prepared rapid gradient echo (MP-RAGE) was used, with the following parameters: repetition time (TR) = 6000 ms, echo time (TE) = 2 ms, flip angle = 12°, number of averages (NA) = 1, inversion time = 1600 ms, voxel size = 270 × 270 × 540 μm, and scan time = 20 min. For T2w imaging, rapid acquisition with relaxation enhancement (RARE) was utilized, with the following parameters: TR = 4000 ms, TE = 22 ms, RARE factor = 4, flip angle = 90°, NA = 1, voxel size = 270 × 270 × 540 μm, and scan time = 7 min, 24 s. For DWI, spin-echo echo-planar imaging was applied, with the following parameters: TR = 3000 ms, TE = 25.6 ms, δ = 6 ms, Δ = 12 ms, b-value = 1000 and 3000 s/mm^2^ in 30 and 60 diffusion directions, respectively (plus 4 b0 images), number of segments = 6, flip angle = 90°, NA = 3, voxel size = 350 × 350 × 700 μm, and scan time = 90 min. Diffusion metrics were created using the diffusion tensor imaging (DTI) model, and the diffusion fiber connectome was created using constrained spherical deconvolution^11^. For rsfMRI, gradient-recalled echo-planar imaging was used, with the following parameters: TR = 1500 ms, TE = 18 ms, number of shots = 1, flip angle = 40°, NA = 1, number of repetitions = 400, voxel size = 500 × 500 × 1000 μm, and scan time = 10 min.

### Treatments of animals in ex vivo imaging

Each animal was perfusion-fixed with 4% paraformaldehyde (PFA), and the brain was dissected out of the cranium and soaked in PFA for ex vivo imaging. During ex vivo imaging, the brain was wrapped in a sponge and soaked in fluorine solution, which exhibits no signal on MRI images, in a plastic container. Vacuum degassing was performed for artifact reduction. The PFA solution used for fixation was replaced with fresh solution weekly to maintain fixation.

### Image acquisition ex vivo

MRI was performed using a 9.4-T BioSpec 94/30 unit (Bruker Optik GmbH) and a transmitting and receiving solenoid type coil with a 28-mm inner diameter. For T2w imaging, RARE was used, with the following parameters: TR = 10,000 ms, TE = 29.3 ms, RARE factor = 4, flip angle = 90°, NA = 16, voxel size = 100 × 100 × 200 μm, and scan time = 3 h, 20 min. For DWI, spin-echo echo-planar imaging was used, with the following parameters: TR = 4000 ms, TE = 28.4 ms, δ = 7 ms, Δ = 14 ms, b-value = 1,000, 3,000, and 5,000 s/mm^2^ in 128 diffusion direction each (plus 6 b0 images), number of segments = 10, flip angle = 90°, NA = 2, voxel size = 200 × 200 × 200 μm, and scan time = 6 h, 39 mins.

### Data processing pipeline

#### Structural image

To correct T2w images, whole brains were extracted from the image data using BrainSuite18a (David W. Shattuck, Ahmanson-Lovelace Brain Mapping Center at the University of California). Mask images were created, and a registration process was performed to align the standard brain images by mapping brain region data to the structural images of each animal. The analysis software ANTs (Brian B. Avants, University of Pennsylvania) was used for this process^12^.

To locate brain area, we digitized the Atlas^13^ proposed by Hashikawa et al.^14^ in 3D setting. Since the Hashikawa atlas was segmented by histology whose resolution scale is extremely high for MRI data analysis, we merged the regional labels into 6 and 52 and 111 anatomically validated regions defined by the anatomist, thereby fitting for both structural and functional MRI analysis.

The migration information from the standard brain image to the structural image of each animal was calculated. This information was superimposed on the brain region data to create information corresponding to the structural image of each animal. The T1w/T2w approach was proposed by Glasser et al. in 2011, and it showed how to increase contrast related to myelin content by calculating a simple ratio between T1w and T2w images^15^. Since this can be calculated from the ratio of T1w and T2w images, there was no need for novel imaging^16^.

#### Diffusion MRI

Pre-processing steps, such as artifact removal, were performed. These processes were conducted using the brain image analysis tool Mrtrix3 version 3.0.3.12 (J-Donald Tournier, School of Biomedical Engineering & Imaging Sciences, King’s College London)^17^. The following commands were used in different processes: dwidenoize, mrdegibbs, dwipreproc, and dwibiascorrect. After image pre-processing, diffusion metrics were created using the DTI model. The diffusion fiber connectome was created using constrained spherical deconvolution (Tournier et al., 2004), and the ex vivo diffusion fiber connectome was created using high angular resolution diffusion-weighted MRI^18^. In this process, we used the MRTrix3 software in tensor analysis and fiber construction (dwi2tensor, tensor2metrics, dwi2response, dwi2fod, taken, and SIFT). We constructed diffusion metric images, axial diffusivity (AD) images, radial diffusivity (RD) images, fractional anisotropy (FA) images, and connectome matrices based on the number of fibers.

#### Resting-state functional image

Data pre-processing was performed using the SPM12 software package (Wellcome Department of Cognitive Neurology, London, UK) running under MATLAB (MathWorks, Natick, MA, USA). We subsequently performed denoising steps with the functional connectivity toolbox (CONN). The empirical blood oxygenation level-dependent signals were band-pass filtered within a narrow band of 0.01–0.08 Hz. The analysis used fMRI data from a 20-min scan (initial 40 volumes discarded; subsequent 560 functional volumes) for awake data and a 10-min scan (initial 20 volumes discarded; subsequent 380 functional volumes) for anesthetized data. The empirical FC matrix was calculated using Pearson correlation between the average time courses of 104 brain regions for 3 awake and 31 anesthetized healthy common marmosets at rest and was averaged across the marmosets.

#### Brain region

In this study, we used the anatomically segmented atlas of the common marmoset brain^11^ reported by Hashikawa et al., and the atlas was applied to the data from 111 regions of one brain and 52 regions of another brain created by combining several regions among 111 regions. The regions of interest (ROIs) are listed in ROImerge_data_v2.xlsx. In addition, the data divided into six regions (cerebrospinal fluid, gray matter, deep gray matter, white matter, cerebellum, and brainstem) were obtained for large segmentation and were fitted to individual brain data. The software ANTs was used to align the brain atlas to the individual brains.

## Data Records

All datasets are publicly available in the Brain/MINDS Data Portal (https://dataportal.brainminds.jp/marmoset-mri-na216). The dataset is divided into four sections, which are as follows: in vivo MRI of 216 animals, ex vivo MRI of 91 animals, standard brain, and BMA 2019 atlas mappings.

The in vivo MRI metadata are described in Individual_information_invivo.xlsx. The metadata include the following information for each of the 216 animals: ID, age, sex, weight, relaxometry image, diffusion MRI image and structural connectome, label image, and anesthesia (as shown in Fig. 1). The in vivo MRI dataset includes the following: T1w image (T1WI_*.nii.gz), T2w image (T2WI_*.nii.gz), myelin contrast image (T1wT2w_*.nii.gz), AD image (dtiAD_*.nii.gz), RD image (dtiRD_*.nii.gz), mean diffusivity (MD) image (dtiMD_*.nii.gz), FA image (dtiFA_*.nii.gz), FA color image (dtiFAc_*.nii.gz), 6 ROI label map image (label006_*.nii.gz), 52 ROI label map image (label052_*.nii.gz), and 111 ROI label map image (label111_*.nii.gz). These can be downloaded in NIFTI format (nii.gz) for each individual marmoset. In addition, the volume for each ROI of the BMA 2019 atlas (Brain_*_summary.xlsx, within i_Variables_gm.zip, with the atlas available at https://doi.org/10.24475/bma.4520), structural connectome between ROIs (DiffusionSC_*.csv), anesthetized functional connectome (AnethFC_*.csv), and awake functional connectome (AwakeFC_*.csv) can be downloaded for each individual.

**Fig. 1:**
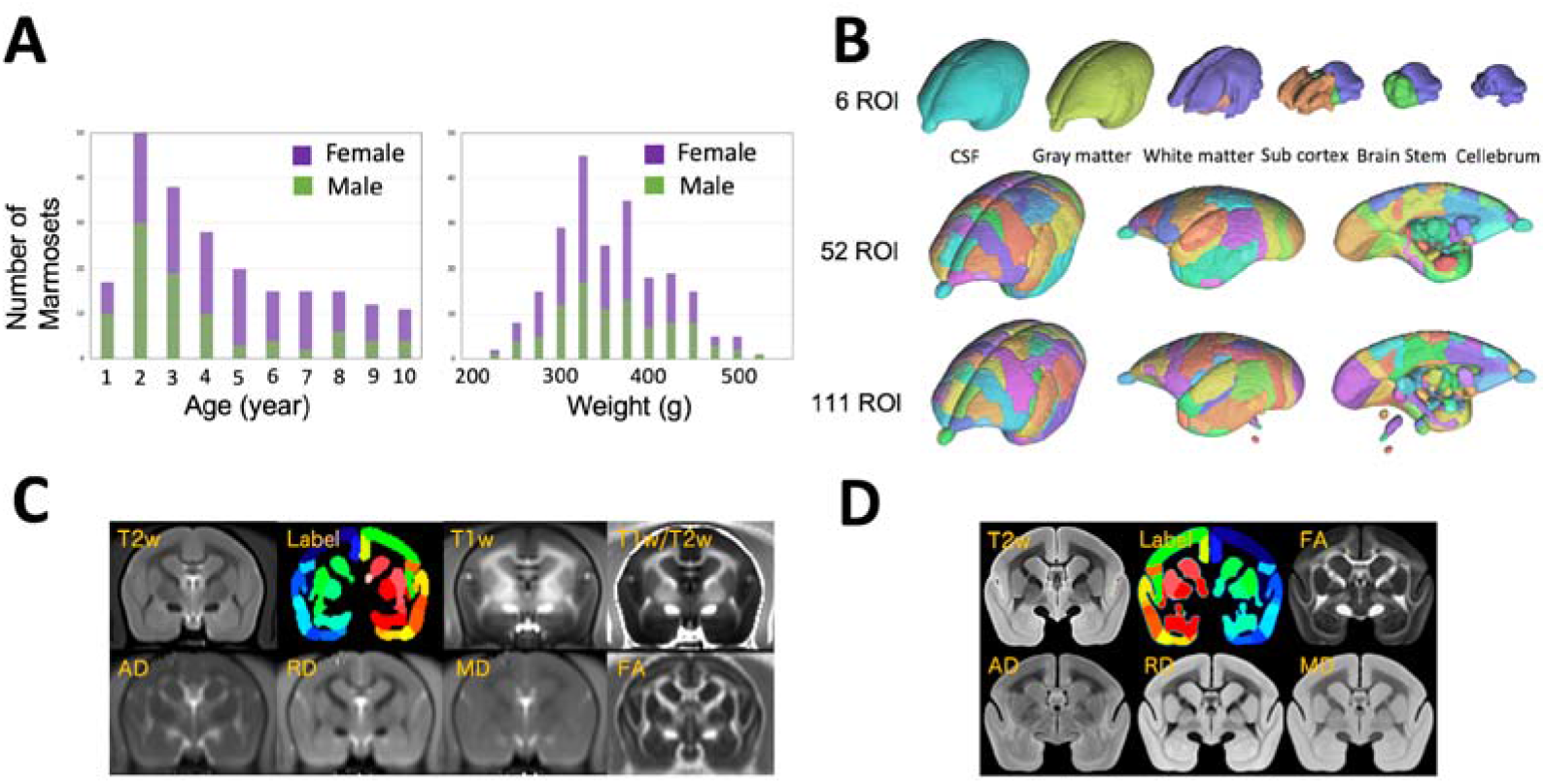
Summary of the magnetic resonance imaging (MRI) dataset of marmosets. A. Histograms of age and weight of marmosets according to sex. B. Multiscale label map with a three-dimensional visualization of merged regions of the Hashikawa atlas into 6, 52, and 111 regions of interest (ROIs). C. For in vivo MRI data, T2-weighted (T2w), label, T1-weighted (T1w), T1w-to-T2w ratio, axial diffusivity (AD), radial diffusivity (RD), mean diffusivity (MD), and fractional anisotropy (FA) images are shown. D. For ex vivo MRI data, T2w, label, FA, AD, RD, and MD images are depicted.

*Ex vivo* MRI data are described in Individual_information_exvivo.xlsx. The metadata include ID, age, sex, weight, T2w image, diffusion MRI image and structural connectome, label image, and ID of the corresponding in vivo MRI data for each of the 91 animals. The ex vivo MRI dataset includes the following: T2w image (T2Wi_ex*.nii.gz), AD image (dtiAD_ex*.nii.gz), RD image (dtiRD_ex*.nii.gz), MD image (dtiMD_ex*.nii.gz), FA image (dtiFA_ex*.nii.gz), FA color image (dtiFAc_ex*.nii.gz), and 52 ROI label map image (label052_ex*.nii.gz). These can be downloaded in NIFTI format for each individual. In addition, the volume of each ROI (Brain_ex*_summary.xlsx, within e_Variables_gm.zip) for the BMA 2019 atlas and the structural connectome (DiffusionSC_*ex*.csv) between the ROIs of each value can be downloaded for each individual.

The standard brain section averaged the images after the registration of the individual data to the BMA 2019 atlas. The BMA 2019 atlas mapping section provides data on the BMA 2019 atlas mapped to the in vivo, ex vivo, and standard spaces of each individual brain.

The section of each dataset can be downloaded as a whole. Moreover, using the dynamic filtering widget (Fig. 2A), it is possible to only display individual data related to certain age groups, sex, and weights, thereby making it easier to limit downloads to a specific part of the dataset. Moreover, most individual data can be previewed online (as shown in Fig. 2B). If different modalities for the same individual are selected, they can be viewed side by side.

**Fig. 2:**
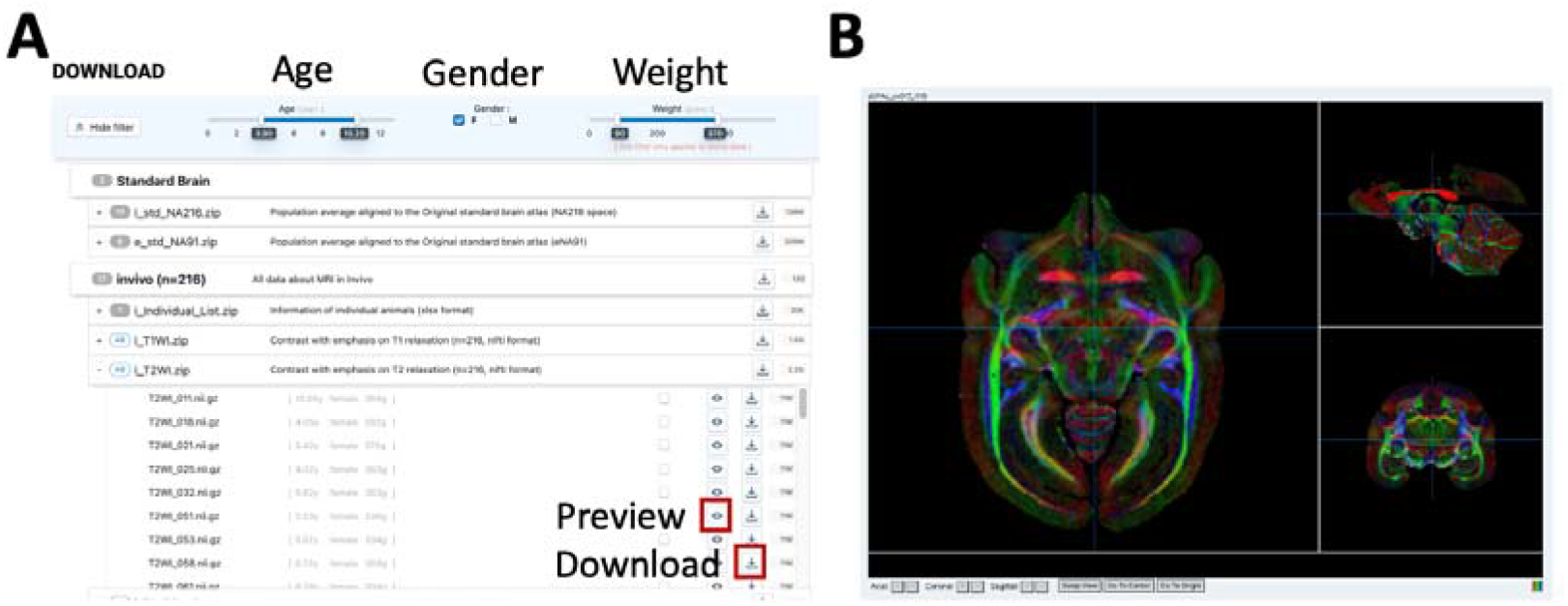
Download page and preview. A. At the top is a filtering widget to display specific age, sex, and weight data. Each file has a Preview and Download buttons. B. Pressing the Preview button opens the individual data viewer, which allows visualizing NIfTI volume slices along the three standard axes.

## Technical Validation

### Brain volume

We evaluated age-related volumetric changes in each brain region by analyzing T2w images, as shown in Figure 3. The volume of the gray and white matter decreased from the developmental age (approximately 12 months) to maturity (approximately 18 months), followed by a gradual downward trend, similar to that observed in humans aged 20–50 years in a similar study^19^. Moreover, a gradual decrease in the cortex volume of common marmosets was consistent with that of a previous study^7^ that assessed the common marmoset brain volume from the age of 1 month to 18 months. Despite the overall reduction in brain volume with age, the proportion of white matter increases with age. The volume of cerebrospinal fluid significantly increases in humans as the brain atrophies with old age^20^. However, no significant increase was observed in this study. Therefore, caution should be exercised for comparative studies of age-related cerebrospinal fluid changes involving humans or other primates. An increase in individual variability was observed with age, and the age-related variance values were 19.27%, 25.18%, and 28.75% for the ages 18– 36 months, 37–72 months, and ≥73 months, respectively. These tendencies are similar to those in humans. Thus, our database can be considered highly reliable and be used for assessing detailed changes in volume with age.

**Fig. 3:**
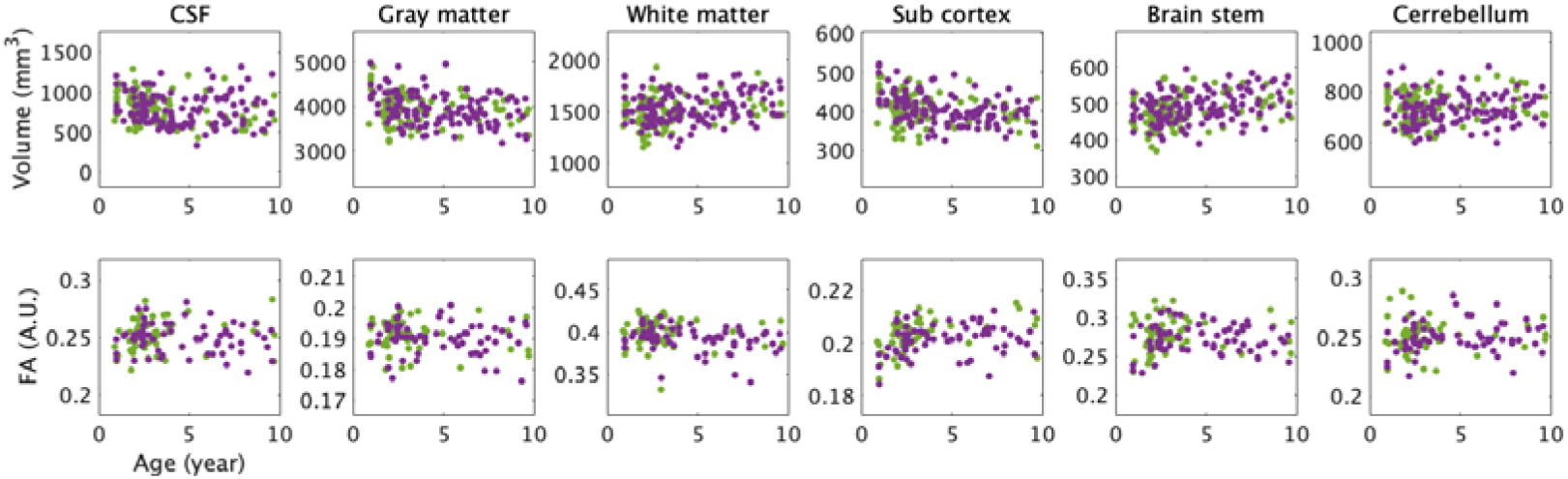
Scatter plot of volume size and mean fractional anisotropy (FA) values versus age for marmosets. The purple points correspond to female marmosets, while the green correspond to male marmosets. The upper panels show the volume of the cerebrospinal fluid (CSF), cortex, white matter, subcortex, brain stem, and cerebellum in relation to age. The lower panels show the FA of the CSF, cortex, white matter, subcortex, brain stem, and cerebellum in relation to age.

### Diffusion structural MRI

The mean values of the DTI metrics including FA, AD, and RDr in each brain region were evaluated and summarized as a scatter plot to confirm data distribution. We observed that FA in the white matter of common marmosets had a slightly higher dispersion than AD and RD in the white matter of marmosets. A large variation in the white matter tissue was observed with different values based on the frontal and posterior areas; data showed a decreasing trend with age. Compared with previous human studies, the FA was consistent with a decrease of up to 40 years of age in humans studies^21^. In the deep gray and white matter, FA increases until 4 years of age, and it does not significantly change thereafter. However, no changes in regions apart from the deep gray matter were observed. In humans, the number increases until 20 years of age, and it does not change significantly thereafter. Next, it declines at approximately 40 years of age^22^. In both common marmosets and humans, the FA is lower than the average FA for all ages until the developmental period. This indicates that myelin development limits intracellular diffusion until development is complete, and FA stabilizes after development. The reason for the decrease in diffusion coefficient with age is that myelination limits proton diffusion in the extracellular space as cells develop^23,24^. The data quality and the diffusion metric analysis are highly reliable.

Figure 4 shows the whole brain connectivity derived via diffusion tractography. Our connectivity matrices in in vivo and ex vivo datasets showed that connections are stronger in the brain network on the ipsilateral side and weaker in the contralateral brain. If the distance between brain regions is longer, the number of connections is fewer in the neural structure network on diffusion MRI. The connection trends between the brain regions are almost similar between the in vivo and ex vivo connections, and the strength was more significant in the *ex vivo* data, which indicated that the time was longer in the *ex vivo* data. Although there are few differences, major diffusion fiber connectivity is observed both in vivo and ex vivo data.

**Fig. 4:**
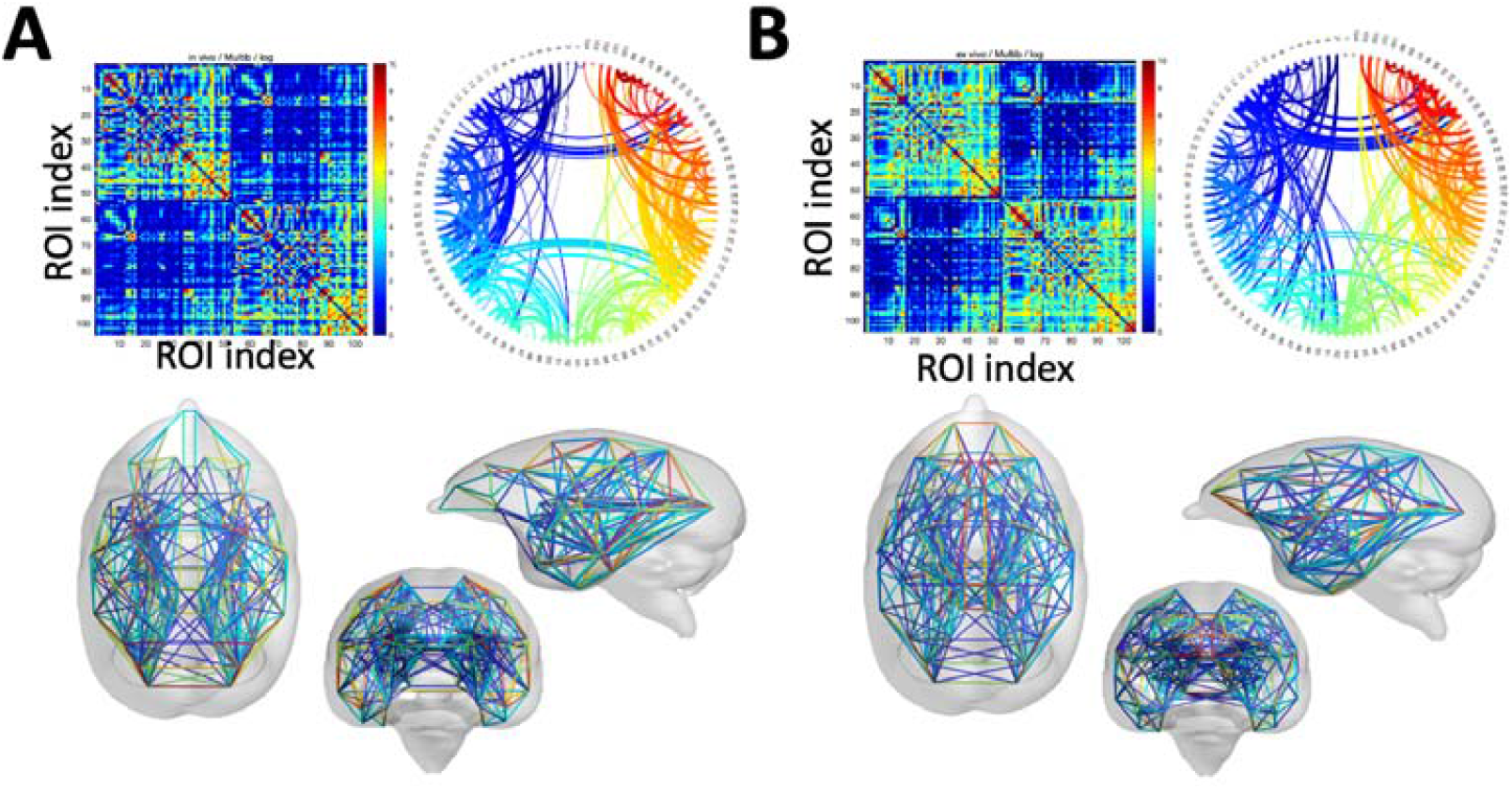
Mean values of structural connectivity estimated from diffusion magnetic resonance imaging (dMRI). A. *In vivo* dMRI: The upper left panel shows the mean of the connectivity matrix for 216 marmosets calculated from tractography. The upper right panel shows the circular connectivity plot with the connections colored based on region, and the lower panel depicts the connectivity plot from the center of gravity of the region of interest (ROI) location according to region. B. Ex vivo dMRI: Upper left panel shows the mean of the connectivity matrix of 91 marmosets calculated from tractography. The upper right panel depicts the circular connectivity plot with the connections colored based on region, and the lower panel shows the connectivity plot from the center of gravity of the ROI location according to region.

### Resting-state functional MRI

To reduce body movement, stress, and pain in animals, most of our rsfMRI data are performed under anesthesia. Under anesthesia, the bold signal of rsfMRI is quite low compared with the awake state, as shown in Figure 5. Hence, the actual brain activities in the resting condition could not be directly evaluated^26^. Recent techniques have been made it possible to evaluate awakefulness rsfMRI available. The current dataset includes data obtained under both anesthesia and awake states. Therefore, the level of brain activity differed between the anesthesia and awake states (as shown in Fig. 5).

**Fig. 5:**
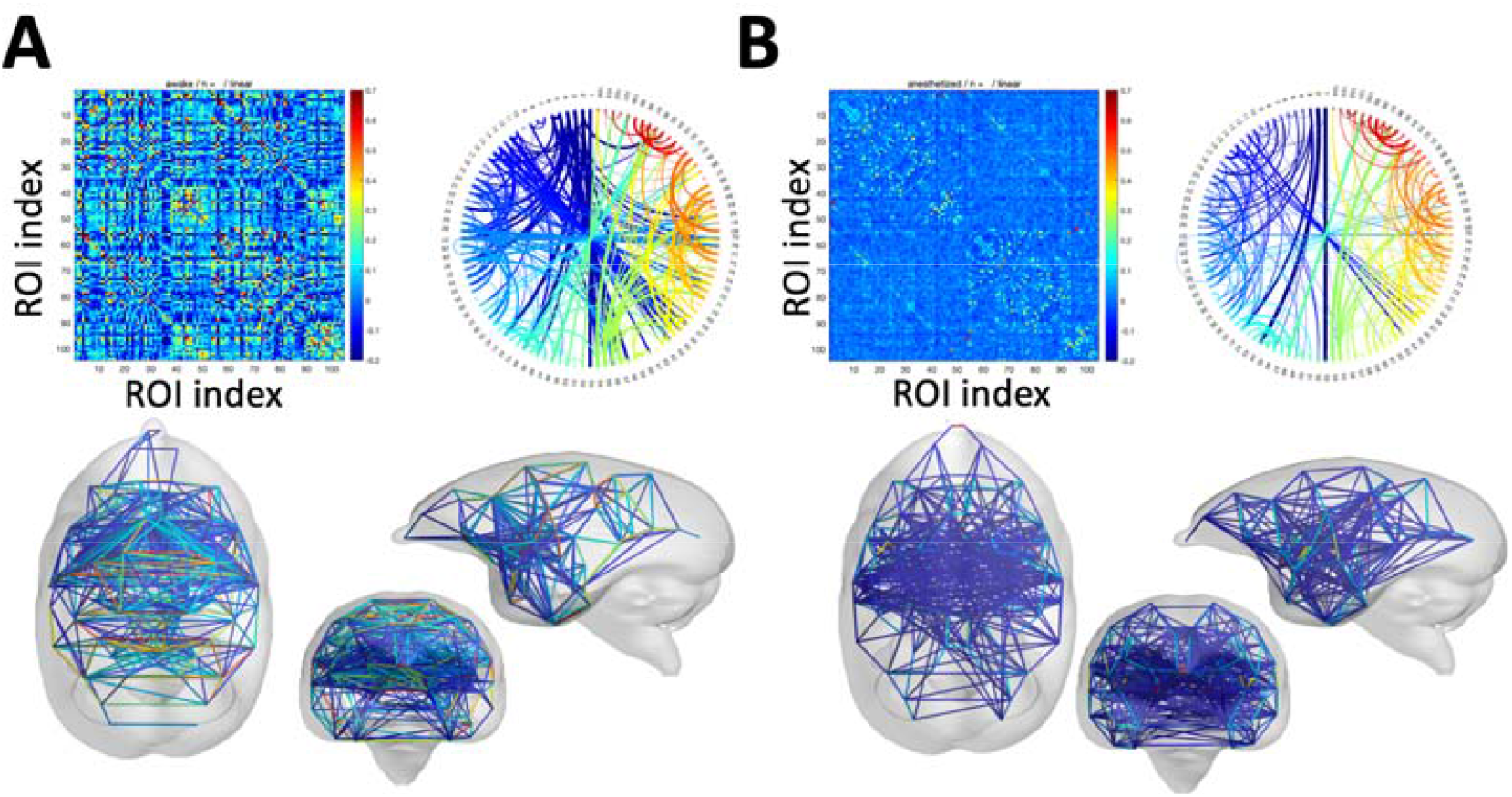
Mean values of functional connectivity estimated from functional magnetic resonance imaging (fMRI). A. Awake fMRI: Upper left panel shows the mean of the cross correlation matrix for three marmosets calculated from fMRI activity. The upper right panel shows the circular connectivity plot with the connections colored according to region, and the lower panel depicts the connectivity plot from the center of gravity of the region of interest (ROI) location based on region. B. Anesthetized fMRI: The upper left panel indicates the mean of the cross-correlation matrix of 31 marmosets calculated from fMRI activity. The upper right panel shows the circular connectivity plot with the connections colored according to region, and the lower panel shows the connectivity plot from the center of gravity of the ROI location based on region.

## Code availability

Brain/MINDS

Data portal https://dataportal.brainminds.jp/marmoset-mri-na216

BrainSuite18a

David W. Shattuck, Ahmanson-Lovelace Brain Mapping Center at the University of California

ANTs (dvanced Normalization Tools)

Brian B. Avants, University of Pennsylvania

Mrtrix3

J-Donald Tournier, School of Biomedical Engineering & Imaging Sciences, King’s College London

SPM12 (Statistical Parametric Mapping package)

Department of Cognitive Neurology, London, UK)

CONN (the functional connectivity toolbox)

Alfonso Nieto-Castanon, Department of Speech, Language, and Hearing Sciences, Boston University

## Acknowledgments

This work was supported by the program for Brain Mapping by Integrated Neurotechnologies for Disease Studies (Brain/MINDS) from the Japan Agency for Medical Research and Development (AMED) (grant number JP15dm0207001 to H.O., JP19dm0207088 to K.N.) and JSPS KAKENHI (grant number JP20H03630 to J.H.), and by “MRI platform” as a program of the Project for Promoting Public Utilization of Advanced Research Infrastructure of the Ministry of Education, Culture, Sports, Science and Technology MEXT, Japan (grant number JPMXS0450400622 to J.H.).

## Author Contributions

Conceptualization: J.H. Methodology: J.H. Software: J.H., K.N., H.T., A.W., Y.H., R.G., M.M., and H.S. Validation: J.H., K.N., H.T., Y.H., M.I., A.U., F.S., N.I., and D.Y. Formal analysis: J.H., K.N., H.T., Y.H., M.I., A.U., F.S., N.I., and D.Y. Investigation: J.H, K.N., H.T., Y.H., M.I., A.U., F.S., N.I., and D.Y. Resources: J.H., K.N., H.T., A.W., H.A., T.T., K.H., N.K., T.S., T.Y., and H.J.O. Data curation: J.H., K.N., H.T., A.W., Y.H., R.G., A.W., H.A., T.T., M.M., H.S., K.H., N.K., T.S., T.Y., and H.J.O. Writing: J.H., K.N., and H.O. Visualization: J.H. Supervision: J.H. and H.O. Project administration: J.H. and H.O. Funding acquisition: J.H., K.N., A.W., T.S., T.Y., H.J.O., and H.O. All authors have read and approved the final manuscript.

## Competing Interests

The authors declare no competing interests.

## Notes

https://dataportal.brainminds.jp/marmoset-mri-na216

### Competing Interest Statement

The authors have declared no competing interest.

